# Human footprint differentially impacts genetic connectivity of four wide-ranging mammals in a fragmented landscape

**DOI:** 10.1101/717777

**Authors:** Prachi Thatte, Anuradha Chandramouli, Abhinav Tyagi, Kaushal Patel, Phulmani Baro, Himanshu Chhattani, Uma Ramakrishnan

**Author notes:** Corresponding author National Centre for Biological Sciences, Tata Institute of Fundamental Research, Bellary Road, Bangalore 560065. /31.

## Abstract

**Aim:** Maintaining connectivity is critical for long-term persistence of wild carnivores in landscapes fragmented due to anthropogenic activity. We examined spatial genetic structure and the impact of landscape features on connectivity in four wide-spread species- jungle cat (Felis chaus), leopard (Panthera pardus), sloth bear (Melursus ursinus) and tiger (Panthera tigris). Location Our study was carried out in the central Indian landscape, a stronghold in terms of distribution and abundance of large mammals. The landscape comprises fragmented forests embedded in a heterogeneous matrix of multiple land-use types.

**Methods:** Microsatellite data from non-invasively sampled individuals (90 jungle cats, 82 leopards, 104 sloth bears and 117 tigers) were used to investigate genetic differentiation. Impact of landscape features on gene flow was inferred using a multi-model landscape resistance optimization approach.

**Results:** All four study species revealed significant isolation by distance (IBD). The correlation between genetic and geographic distance was significant only over a short distance for jungle cat, followed by longer distances for sloth bear, leopard and tiger. Overall, human footprint had a high negative impact on geneflow in tigers, followed by leopards, sloth bears and the least on jungle cats. Individual landscape variables- land-use, human population density, density of linear features and roads- impacted the study species differently. Although land-use was found to be an important variable explaining genetic connectivity for all four species, the amount of variation explained, the optimum spatial resolution and the resistance offered by different land-use classes varied.

**Main conclusions:** As expected from theory, but rarely demonstrated using empirical data, the pattern of spatial autocorrelation of genetic variation scaled with dispersal ability and density of the study species. Landscape genetic analyses revealed species-specific impact of landscape features and provided insights into interactions between species biology and landscape structure. Our results emphasize the need for incorporating functional connectivity data from multiple species for landscape-level conservation planning.

## Introduction

Habitat destruction and fragmentation due to human footprint is degrading habitat quality and increasing extinction risk of mammals globally (Crooks et al., 2017). Dispersal and genetic exchange between habitat fragments is critical for reducing extinction risk, maintaining genetic diversity and persistence of sub-divided populations (Moilanen & Nieminen, 2002; Thatte et al., 2018a). Maintaining connectivity is hence recognized as a key factor in conservation and management of endangered mammal species.

It is no surprise therefore that research on connectivity conservation has markedly increased over the last decade (Correa Ayram et al., 2016). However, connectivity studies in mammals have largely focused on a single focal species, usually a large predator (Beier et al., 2008; Segelbacher et al., 2010; McRae et al., 2012; Dudaniec et al., 2013; Joshi et al., 2013; execept in a few recent cases- Dudaniec et al., 2016; Wultsch et al., 2016; Marrotte et al., 2017). Conservation strategies based on information from a single species may not effectively capture varied ecological requirements for dispersal of other sympatric species (Brodie et al., 2015; Gangadharan et al., 2016).

Co-occurring species may respond differently to fragmentation based on the differences in body size, dispersal ability, generation time, area requirement and distribution (Gehring & Swihart, 2003; Davidson et al., 2009). Landscape structure or configuration of habitat patches in a landscape may also determine response to fragmentation. Habitat association and the ability to move through heterogeneous matrix between habitat patches can further affect species response to fragmentation (Kierepka et al., 2016). In summary, connectivity for different species may vary based on their biology, landscape structure and interaction between the two, even in the context of a single landscape.

Conservation strategies for a landscape need to factor in the different requirements of species to prioritize areas of maximum diversity as well as areas where the most endangered species are likely to persist in the long term. For long-term persistence, areas that facilitate dispersal of multiple species need to be protected. While opinion articles emphasize the importance of multi-species landscape genetics studies when developing guidelines for landscape level conservation and management (Keller et al., 2015; Richardson et al., 2016), relatively few studies have compared connectivity for multiple species (eg. Engler et al., 2014; Dudaniec et al., 2016; Wultsch et al., 2016; Marrotte et al., 2017).

We studied genetic connectivity in four wide-spread species- jungle cat (Felis chaus), leopard (Panthera pardus), sloth bear (Melursus ursinus) and tiger (Panthera tigris) in the Central Indian landscape. Classified as a global priority tiger conservation landscape, it is a stronghold in terms of mammal distribution and abundance (Jhala, Y. V.;Qureshi, Q;Gopal 2015). Several studies have investigated tiger connectivity in the landscape (Joshi et al., 2013; Sharma et al., 2013; Yumnam et al., 2014; Reddy et al., 2017; Thatte et al., 2018a). However, there is dearth of information on habitat and population connectivity for other sympatric species (Dutta et al., 2013a, 2015).

Our study species vary in body size, dispersal ability, distribution and habitat association, traits that determine species sensitivity to habitat fragmentation (Davidson et al., 2009; Kierepka et al., 2016). Interestingly, considerations based on the above factors result in contradictory predictions in terms of expected spatial genetic structure for our study species (discussed below and represented in Figure 1).

**Figure 1.**
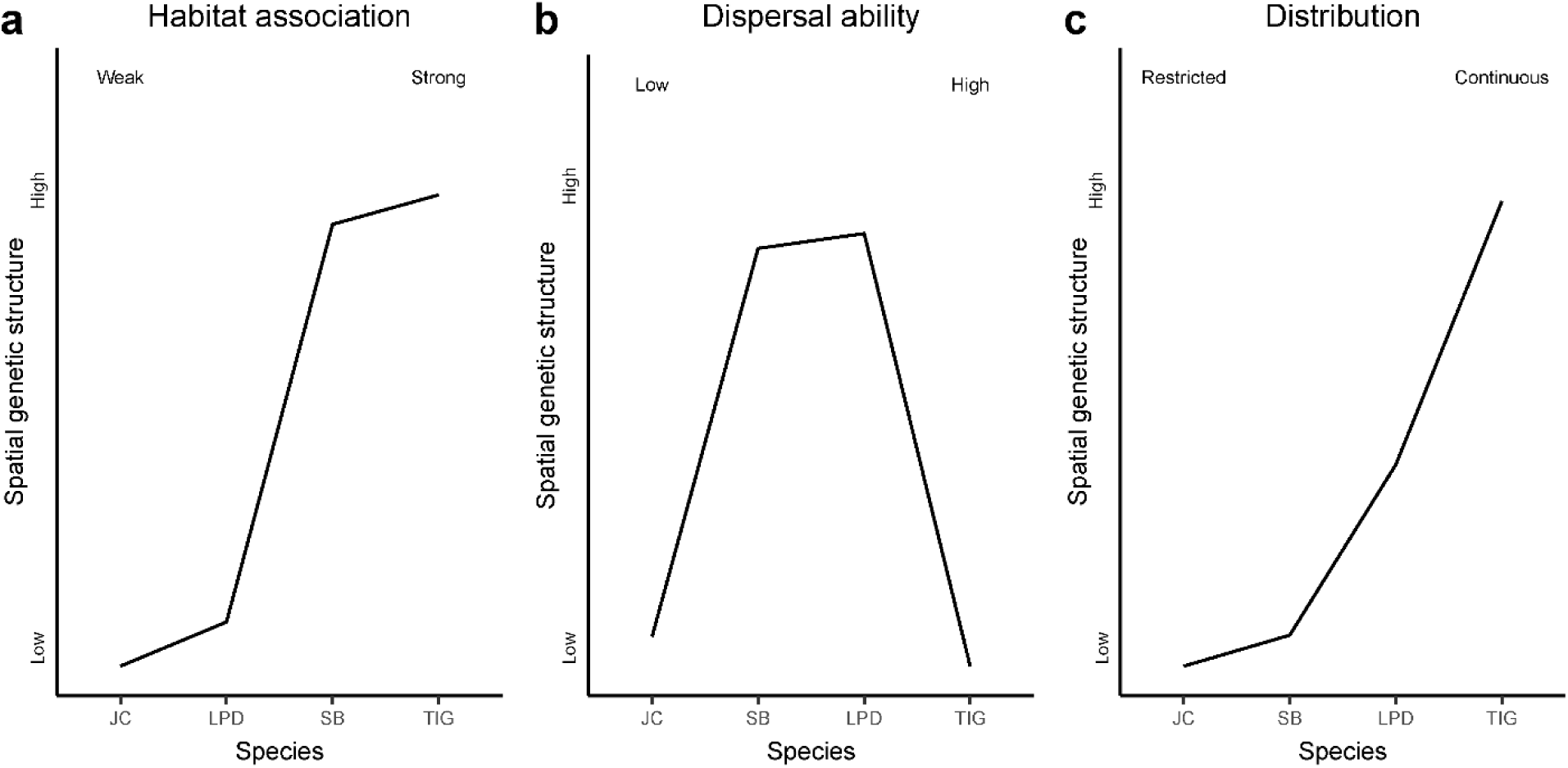
Spatial genetic structure expectations based on habitat association, dispersal ability and distributions of the study species.

Empirical evidence across taxa suggests that organisms with specific habitat requirements (and hence habitat association) are usually characterized by low gene flow resulting in strong genetic differentiation (Stuart-Fox et al., 2001; Zayed et al., 2006; Khimoun et al., 2016; Kierepka et al., 2016; Harvey et al., 2017). Among our study species, tigers and sloth bears show stronger habitat associations (Kitchener & Dugmore, 2000; Ramesh et al., 2012; Joshi et al., 2013; Das et al., 2014; Yumnam et al., 2014; Puri et al., 2015) and hence we expect them to have higher genetic differentiation than jungle cats and leopards (Athreya et al., 2013; Gray et al., 2016; Stein et al., 2016).

Dispersal ability of a species can also impact genetic structure. High dispersal ability of a species is likely to homogenize genetic variation over space. Although species with low dispersal ability are likely to have low geneflow, their abundance in a given area is likely to be high (Jenkins et al., 2007; White et al., 2007), resulting in low impact of genetic drift and hence low genetic differentiation. Species with moderate dispersal ability may show higher genetic differentiation: geneflow may not be high enough to homogenize genetic differentiation and the abundance may not be high enough to minimize differentiation. Among our study species, median dispersal distance (calculated based on body size using allometric scaling equations (Sutherland et al., 2000; Bowman et al., 2002) is lowest for jungle cat (8 km), followed by sloth bear (13 km), leopard (35 km) and tiger (85 km) among the study species. Hence we expect sloth bear and leopard to have higher genetic differentiation than tiger and jungle cat.

Continuous and uniform population distribution has been shown to result in lower genetic differentiation (Cowled et al., 2009; Llorens et al., 2017). Spatial patchiness can restrict gene flow and hence increase genetic differentiation. Sloth bears are the most widely distributed large carnivores in central India, followed by leopards and tigers (Jhala et al., 2015). Hence, we expect tigers to have highest genetic differentiation followed by leopard and sloth bear. Jungle cats are also likely to be widely distributed in central India (Gray et al., 2016) although no research studies have been carried out. Hence we expect low genetic differentiation for jungle cats as well.

In summary, all three factors (habitat association, dispersal ability and distribution) predict jungle cats to have the lowest spatial genetic structure among the study species. Predictions for the other three study species are inconsistent. The above predictions ignore the potentially differential ability of the study species to navigate through human modified landscape. In this paper, we seek to address the genetic structure predictions based on dispersal ability, habitat association and distribution with empirical data on four species and also investigate how landscape features impact their movement.

## Methods

### Study area and sampling

With 34% forest cover, the central Indian landscape (CIL) is a stronghold in terms of mammal distribution and abundance (Yoganand et al., 2006; Jhala et al., 2011). 8.5 % of the forested area (∼3 % of the total study area) is protected. Despite being a global hotspot of carnivore habitat, the central Indian landscape has been changing rapidly (Venter et al., 2016).

Similar to other tropical landscapes in the Americas, Asia and Africa, central India faces the threat of further habitat degradation due to anthropogenic activities (Crooks et al., 2011). Being a mineral-rich area, patches of forest have been cleared for mining and the landscape is dotted with mines. There are several railway lines and national and state highways crisscrossing through the landscape. There are five major urban centers (with a population of >1 million) and several big towns. Villages surrounded by agricultural fields cover ∼64% of the land area.

We collected non-invasive scat samples (n=1,025), between November 2012 and April 2017, from forested areas inside and outside protected areas. Existing roads/trails were used to search for potential tiger, sloth bear, leopard and jungle cat scats and fresh scats were collected. Each road/trail was sampled only once so as to reduce recaptures and maximize the area covered.

We sampled nine protected areas: out of which (1) Kanha Tiger Reserve (KTR), (2) Pench Tiger Reserve (PTR), (3) Bandhavgarh Tiger Reserve (BTR), (4) Achanakmar Tiger Reserve (ATR), (5) Nagzira-Nawegaon Tiger Reserve(NN), (6) Satpura Tiger Reserve (STR), (7) Sitanadi-Udanti Tiger Reserve (S-U) and (8)Panna Tiger Reserve (PAN) are International Union for the Conservation of Nature (IUCN) category II protected areas and (9) Tipeshwar Wildlife Sanctuary (TIP) which is a IUCN category IV PA. Nine territorial forest divisions outside these protected areas that fall under IUCN categories IV and VI: (1) North and South Balaghat Forest Divisions (BAL), (2) Bramhapuri Forest Division (BPR), (3) North and South Gondia Forest Divisions (GON), (4) Bhandara Forest Division (BHA), (5) Kawardha Forest Division (KAW), (6) Khairagarh Forest Division (KHA) and (7) South Seoni Forest Division (SEO) were also sampled. Tiger samples collected and used in this study were also a part of Thatte et al. (2018).The scat samples collected between 2012 and 2015 were stored in 30 ml wide mouth bottles containing absolute alcohol. During 2016-17, samples were collected using swabs (HiMedia Inc.) and stored in vials containing Longmire’s buffer (Longmire et al., 1997).

### Genotyping and population genetic analysis

In order to quantify genetic connectivity, we first extracted DNA from scat samples using QIAamp DNA Stool Mini Kit (QIAGEN Inc.) and HiPurA Stool DNA Purification Kit (HiMedia, India). We then identified species using published primers for tiger, jungle cat and leopard and a species specific primer that was designed for sloth bear (Farrell et al., 2000; Thatte et al., 2018b). Jungle cat positive samples were genotyped using 14 felid microsatellite loci (FCA126, FCA069, FCA672, FCA304, FCA628, FCA279, FCA232, msHDZ170, msFCA506, msFCA453, msF42, msFCA441, msFCA391, msF41). For leopards the same set of loci were used, except that msF115 was used instead of msF41. For sloth bears, a panel of 13 bear microsatellite loci was used (G10H, G10X, G10J, G10C, G10O, UarMu59, UarMu05, UarMu09, UarMu23, UarMu51, UarMu61, UarMu10 and UarMu64). G10O and UarMu10 were found to be homozygous for all the samples and hence were not used for analysis. Genotyping details for tigers are reported in Thatte et al., (2018a). To minimize the impact of genotyping error, the genotyping procedure was repeated four times (4 independent PCRs) for each sample, for each locus (Mondol et al., 2009). Genotyping error rates were calculated using GIMLET (Valière, 2002). Individual identification was conducted using the identity analysis module in the program CERVUS (Marshall et al., 1998). Further, heterozygosity based differentiation statistics were calculated using PopGenReport (Adamack & Gruber, 2014), MMOD (Winter, 2012), and HIERFSTAT (Goudet, 2005) in R (Ihaka & Gentleman, 1996). Inter-individual genetic distances was calculated based on proportion of shared alleles (D_PS_; (Bowcock et al., 1994)) using the package ‘adegenet’ (Jombart, 2008) in R.

### Isolation by distance and spatial autocorrelation

Isolation by distance (IBD), the correlation between genetic (D_PS_) and geographic (Euclidean) distance, was calculated using ‘ecodist’ (Goslee & Urban, 2007) package in R. Spatial autocorrelation analysis was carried out using GenAlEx version 6.5 (Peakall & Smouse, 2012). Genetic and geographic pairwise distance matrices were correlated for a set of distance classes (5 km-125 km for jungle cat and 20 km-500 km for the other three species). The correlation coefficient (r) obtained for each distance class was plotted as a correlogram. The significance of r was tested against a null hypothesis (r=0) by generating 95% confidence interval for each distance class (999 permutations, 99 bootstrap repeats).

### Landscape genetics analysis

The relationship between the observed genetic structure and the landscape variables likely to affect gene flow in the study species was systematically evaluated using a multi-model inference and optimization approach (based on Shirk et al., 2010) described in the following sections.

### Landscape variables

We used four landscape variables to build resistance models for all the species- (1) land-use land-cover, (2) human population density, (3) roads and (4) density of linear features (Figure S1). We used Bhuvan (the geo-platform of the Indian Space Research Organization) land-cover data (http://bhuvan.nrsc.gov.in/bhuvan_links.php) and reclassified it into four broad land-cover types which were ranked in order of increasing resistance: (i) forest- this included deciduous and evergreen forest. Central India is dominated by dry and moist deciduous forests and hence the study area hardly has any pixels classified as evergreen, (ii) Scrub and degraded forest-this category also included orchards and plantations, (iii) agriculture-includes zaid, rabi, kharif croplands, those with two or three crops every year, also fallow and wasteland and (iv) built-up areas- areas of human habitation that is non-agricultural and has buildings, roads and other infrastructure. The rank order of the categories was based on previous research (Kitchener & Dugmore, 2000; Joshi et al., 2013; Yumnam et al., 2014). A layer of human population density, developed at the Foundation for Ecological Research, Advocacy and Learning (FERAL), based on the 2011 census of India data was used for this study. We used population density layer as a continuous variable. A vector layer of national highways, state highways and major public roads was reclassified into 6 categories based on the intensity of traffic on the road (based on Passenger Car Unit (PCU) data for 2006 from the Ministry of Road Transport and Highways). PCU is a measure of the impact of a vehicle on traffic and is calculated by converting vehicle units of different types of vehicles (e.g., trucks, buses, motorcycles) into passenger car units. For example, a motorcycle is equivalent to 0.5 passenger cars, whereas a bus is 3.5 PCUs. Road width (1, 2, 4 or 6 lanes) data was used wherever PCU data was not available. The following criteria was used for classification of roads (Indian Roads Congress, 2007, 2010): minor roads (PCU< 1500), very low traffic roads (2500- 5000 PCU- single lane roads), low traffic roads (5000- 10000 PCU- intermediate lane roads), moderate traffic roads (10000- 25000 PCU- 2 lane roads), high traffic roads (25000- 55000 PCU- 4 lane roads), very high traffic roads (>55000 PCU- 6 lane roads). In order to calculate the density of linear features, road, railway lines and irrigation canal vector layers were combined into a single poly-lines layer. This layer was developed at Foundation for Ecological Research, Advocacy and Learning. The density of lines in each cell was calculated to generate a raster layer that was used for analysis.

These four landscape variables were used to carry out multi-scale landscape genetics modelling (described in the next section). Studies have shown that the spatial scale at which a landscape variable is measured can affect the species-landscape relationship (Jackson & Fahrig, 2015). The scale at which a species responds to the surrounding is unknown. The common approach for dealing with the scale problem is analyzing the landscape variables at multiple scales using varying ecological neighborhoods (Krishnamurthy et al., 2016; Zeller et al., 2016) or at varying resolutions (Zeller et al., 2012). The two continuous variables, density of linear features and human population density, were analysed at multiple scales or ecological neighborhoods (0.25 km, 0.5 km, 1 km, 2.5 km, 5 km, 10 km, 25 km and 50 km with a pixel size of 0.25 km) weighted by a Gaussian kernel using the ‘kernel2dsmooth’ function from the smoothie package in R (Gilleland, 2013). These ecological neighborhoods ranged from less than the radius of the home range to more than the median dispersal distance (as suggested by (Jackson & Fahrig, 2015)), except for the tiger for which we did not have a neighborhood size larger than its median dispersal distance. Allometric scaling equations (Bowman et al., 2002) based on body size and trophic level predict the radius of home range and median dispersal distance in the study species to be 0.6 km and 8 km for jungle cat, 1.1 km and 13 km for sloth bear, 2.9 and 35 km for leopard and 6.9 km and 85 km for tiger respectively. LULC, a categorical variable, was analysed at 6 different spatial resolutions (0.25 km, 0.5 km, 1 km, 2.5 km, 5 km, 10 km).

### Multi-model optimization

We used a multi-model optimization approach to identify which landscape variables offer resistance to species movement, the magnitude of resistance (Rmax) for each contributing variable and the functional form or shape of the relationship between each variable and resistance (x). Genetic data was used as a response variable to infer the landscape resistance values and the functional form of the relationship between each variable and resistance. For each landscape variable, we explored a range of parameter values for Rmax and x and identified which combination of values best explained the genetic data. For each variable, six levels of maximum resistance (Rmax = 2, 5, 10, 50, 100, 1000) and six levels of the shape parameter (x= 0.01, 0.1, 1, 2, 5, 10, Figure S2 represents the transformations) were evaluated. Shape parameter controlled the functional response between the landscape variable and resistance to gene flow– 1 meant a linear relationship while values less than and greater than 1 were non-linear. For each combination of Rmax and x, cost-distance between pairs of individuals was calculated. Cost-distance is the modified Euclidean distance between two points that accounts for the resistance offered by the landscape. Linear mixed effects (LME) model with a maximum likelihood population effects (MLPE) parameterization (Clarke et al., 2002) fitted to the data using MLPE.lmm function from the ResistanceGA package (Peterman, 2018) was used to relate genetic distance and cost distance. The best fitting model was identified as the one with the lowest AIC score from the MLPE mixed effects model for each landscape variable.

For each landscape variable, we identified the scale and the combination of parameters (Rmax and x) of the best fitting model. The best fitting univariate models identified based on AIC values for each landscape variable were combined (additively) and optimized again in a multivariate context to account for interactions between different landscape variables for all the study species (Shirk et al., 2010). For multivariate optimization, we varied the model parameters (Rmax and x) of one landscape variable at a time, while holding the parameter values of the other variables constant. If the optimal parameters changed for a landscape variable in the multivariate context, we held the new model parameters constant and varied the parameters of the next variable. We repeated this process until the parameterization of all the variables stabilized.

## Results

### Population genetic analysis

Out of the collected scat samples, based on genetic identification in the lab, 298 were tigers, 343 were sloth bears, 165 were jungle cats and 138 leopards. The remaining 81 samples either failed to amplify with any of the primers used or were from non-target species. Based on unique genotypes, we identified 117 tigers, 104 sloth bears, 92 jungle cats and 82 leopards (Figure 2). 116 of the 117 tiger individuals were also a part of Thatte et al. (2018a). Genotyping error rate, expected and observed heterozygosity (H_e_ and H_o_ respectively) and the probability of identity (P(ID)) are presented in Table 1. The P(ID) (the probability of two different individuals having the same genotype) and the more conservative measure Sib P(ID) (PID when all individuals in the population are assumed to be siblings) was low for all the species, indicating that even related individuals would have a very low probability of having identical genotypes. Sloth bear had lower heterozygosity and higher P(ID) among the study species. The genetic differentiation, estimated as F_ST_ and G_ST_ followed a pattern that was expected based on the spatial distribution of the species (Figure 3).

**Figure 2.**
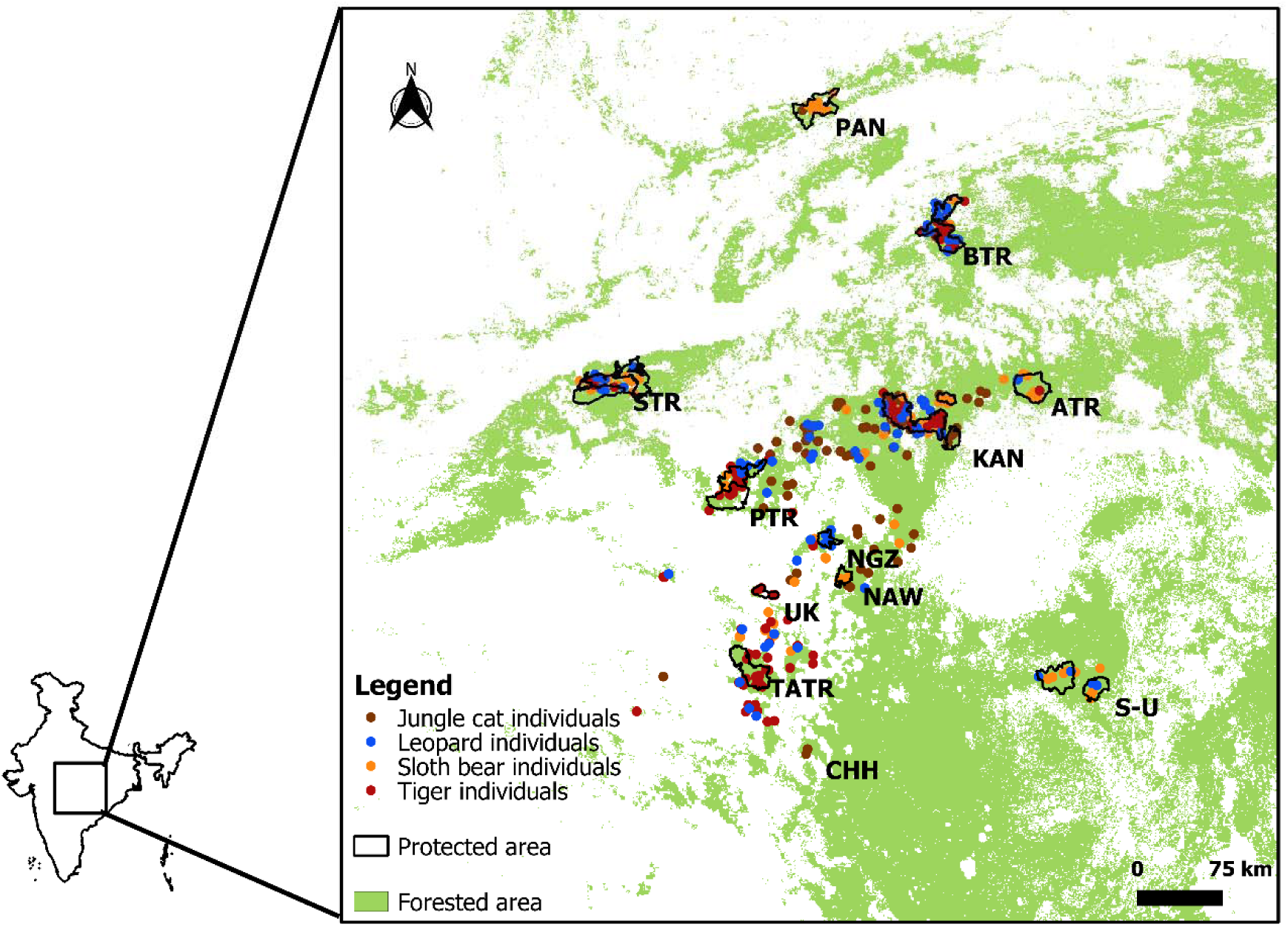
Study area and sampling. The figure contains a map of India with the study landscape highlighted and enlarged. In the enlarged study area, protected areas are marked by black outline and genetically identified individuals of study species as coloured dots.

**Figure 3.**
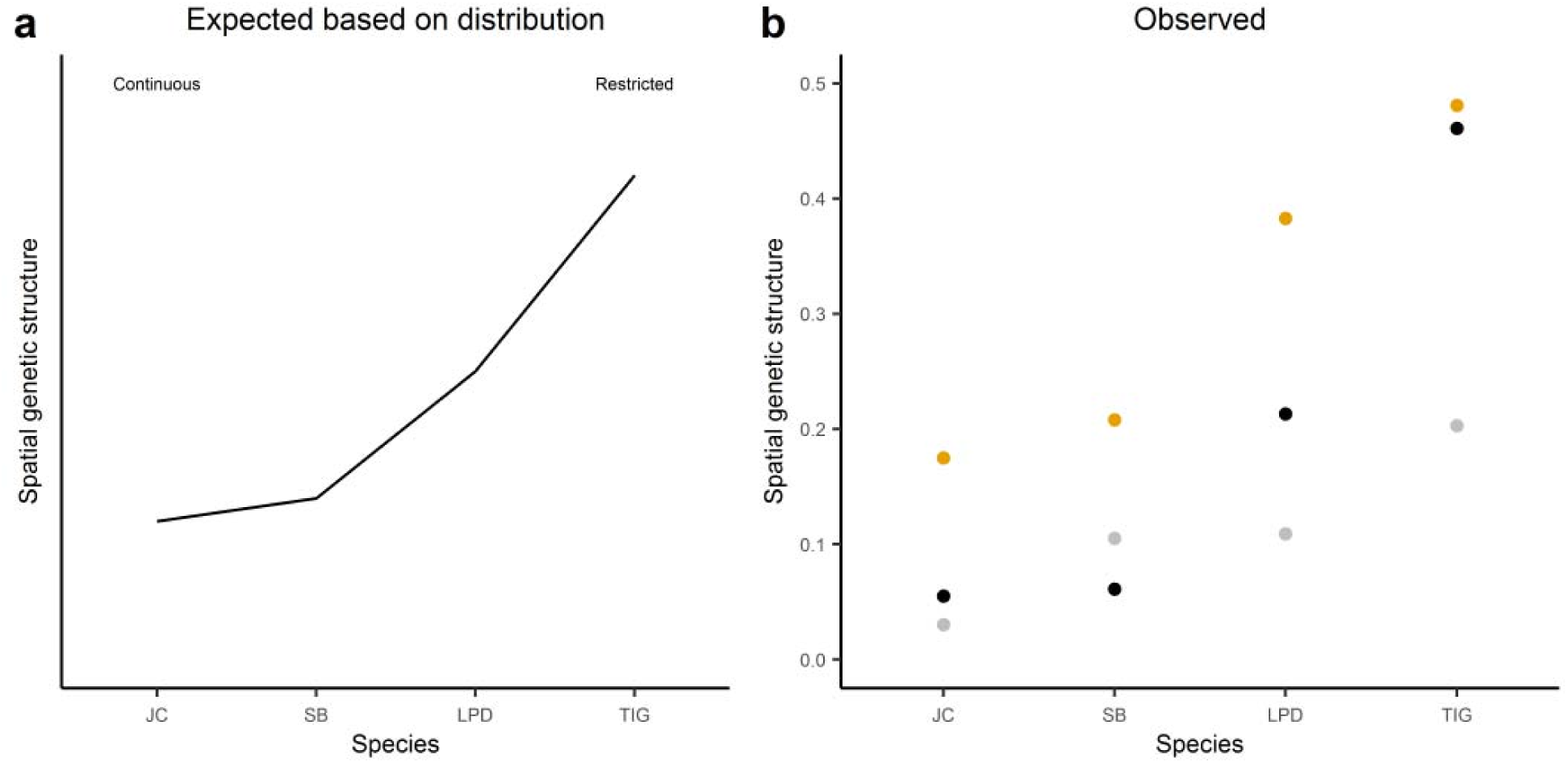
Observed genetic differentiation matched the genetic differentiation pattern expected based on distribution.

**Table 1.** Genotyping error and allele frequency based statistics. He and Ho represent the expected and observed heterozygosities, respectively. P(ID) represents the probability of two different individuals having the same genotype and Sib P(ID) is the PID when all individuals in the population are assumed to be siblings.

| Species | Allelic Dropout | False alleles | He | Ho | P(ID) | Sib P(ID) |
| --- | --- | --- | --- | --- | --- | --- |
| Jungle cat<br>(N= 90) | 0.045 | 0.045 | 0.794 | 0.688 | $5.12 \times 10^{-18}$ | $0.92 \times 10^{-6}$ |
| Leopard<br>(N= 82) | 0.049 | 0.018 | 0.778 | 0.607 | $4 \times 10^{-18}$ | $4.1 \times 10^{-7}$ |
| Sloth bear<br>(N= 104) | 0.060 | 0.030 | 0.51 | 0.392 | $1.6 \times 10^{-7}$ | $1.4 \times 10^{-3}$ |
| Tiger<br>(N= 117) | 0.062 | 0.019 | 0.723 | 0.524 | $1.48 \times 10^{-11}$ | $5.1 \times 10^{-5}$ |

### Isolation by distance

All four study species displayed significant correlation between genetic and geographic distance (IBD for jungle cats r=0.055, p=0.059; for leopards r=0.213, p=0.001; for sloth bears r=0.061, p=0.056 and for tigers r=0.461, p=0.001). To examine the spatial extent of genetic structure, spatial autocorrelation analysis was carried out. IBD broke down at different distances for different species. IBD relationship broke down at distances beyond ∼15 km for jungle cat, followed by ∼40 km for sloth bear, ∼60 km for leopard and ∼140 km for tiger. Figure 4 presents the correlograms showing correlation between genetic and geographic distance (r) as a function of geographic distance classes.

**Figure 4.**
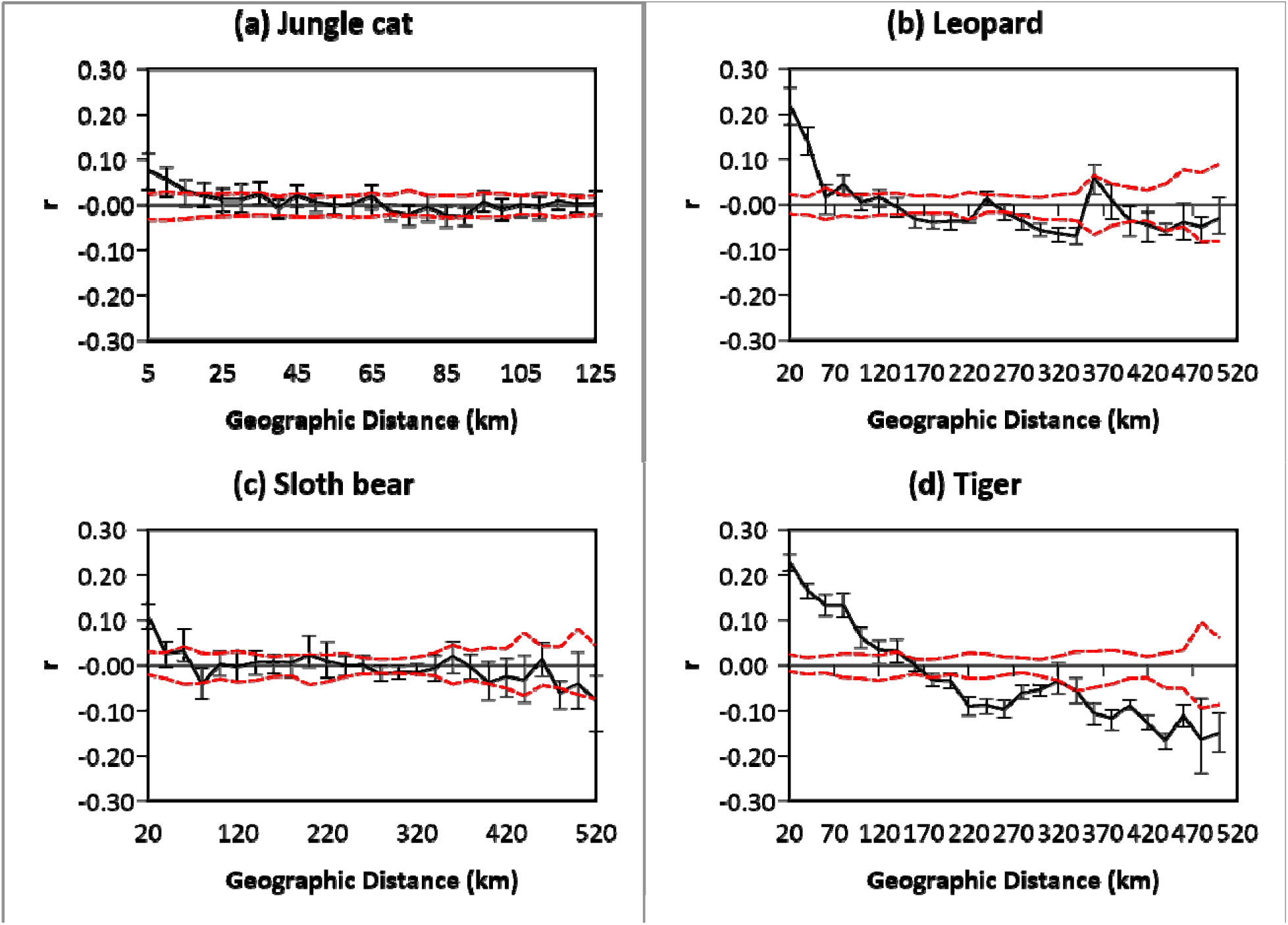
Spatial autocorrelation. x-axis represents distance classes in km. y-axis is the correlation coefficient for the correlation between genetic and geographic distance for samples falling within each distance class. Jungle cats and sloth bears show low correlation (r) and the relationship breaks down at ∼20-25 km and ∼30-60 km respectively. Leopards and Tigers show higher correlation and the relationship breaks down ∼60 km and 140-180 km respectively.

### Landscape genetic analysis

Overall among the study species, landscape variables were found to have the highest impact (based on model fit) on tigers, followed by leopards, sloth bears and jungle cats. Land-use land-cover (LULC) was found to be the most important variable explaining genetic connectivity of all the study species, except jungle cat where it was the second most important variable. However, the resistance offered by different land-use classes, strength of correlation and the optimum spatial resolution varied.

For jungle cat, the effect of landscape variables on genetic connectivity was very low (Table 2). Despite the low effect, we combined the four variables (density of linear features, traffic intensity on roads, human population density and LULC) additively for multivariate optimization. On combining, the model fit increased marginally and the maximum resistance offered by the landscape variables (R_max_) and the shape of the relationship between the landscape variable and the resistance offered (x) changed (Table 3). Human population density, on combining the layers, was found to offer no resistance to jungle cat movement and only built-up areas offered resistance among the land-use classes. Density of linear features was found to have a high negative impact. Despite the low model fit, the optimum model parameters were not sensitive to the model selection method (optimization was also carried out using partial Mantel’s test. Data not presented). Figure 5 presents the relationship between the landscape variables and the resistance they offer and Figure 6 presents the optimized resistance surface for jungle cats.

**Figure 5.**
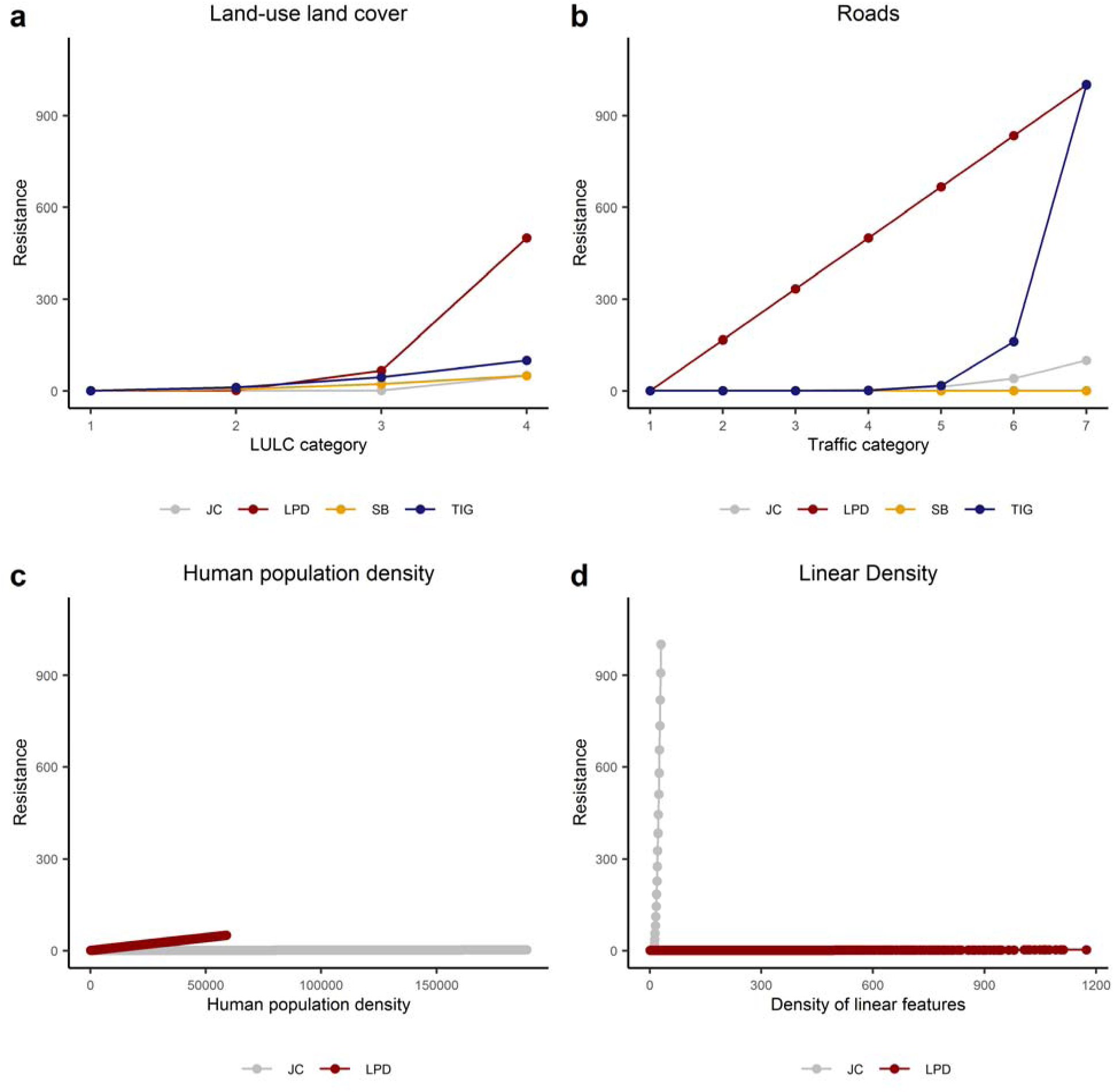
Relationship of landscape features with resistance. x-axis represents each land-use variable used. The categories for roads represent 1- no roads, 2- minor roads, 3- very low traffic roads, 4- low traffic roads, 5- moderate traffic roads, 6- high traffic roads and 7- very high traffic roads. Land-use land-cover categories: 1- forest, 2- degraded forest and scrub, 3- agriculture and 4- built-up. Density of linear features and human population density, being continuous variables, had different maximum values at different resolutions (scales). The optimum scales for different species are different. y-axis is the resistance offered by the different categories/values of the landscape variable.

**Figure 6.**
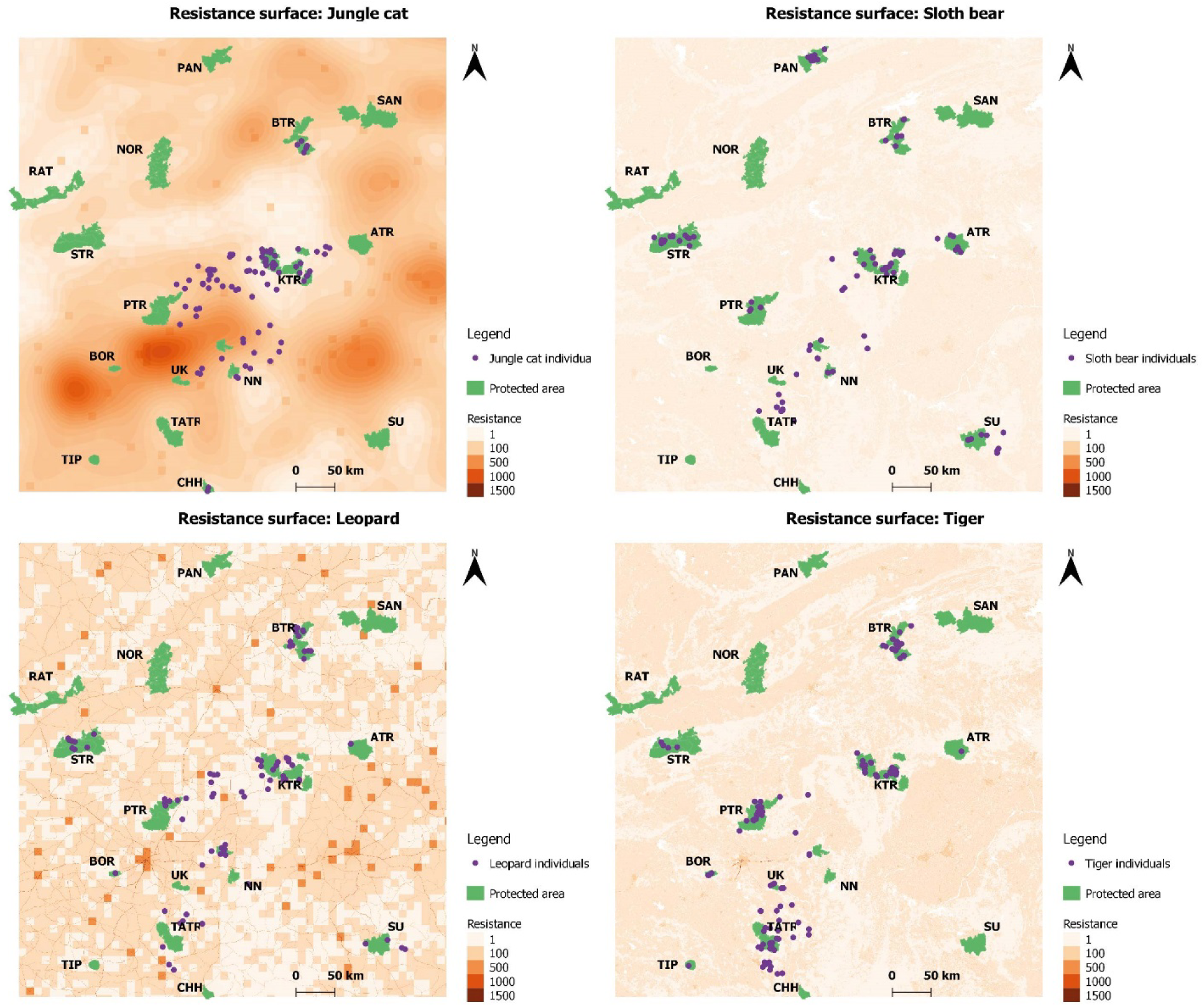
Optimized resistance surface. Dark colours represent high resistance and light colours represent low resistance. Purple dots in each map are the individual locations and green polygons are protected areas. RAT-Ratapani wildlife sanctuary, NOR- Noradehi wildlife sanctuary, BTR- Bandhavgarh tiger reserve, SAN- Sanjay tiger reserve, PAN- Panna tiger reserve, STR- Satpura tiger reserve, ATR- Achanakmar tiger reserve, KTR- Kanha tiger reserve, PTR- Pench tiger reserve, NN- Nagzira Navegaon tiger reserve, BOR- Bor tiger reserve, UK- Umred Karhandla wildlife sanctuary, TATR- Tadoba Andhari tiger reserve, TIP- Tipeshwar tiger reserve, CHH- Chhaprala wildlife sanctuary and SU- Sitanadi Udanti tiger reserve

**Table 2.** Univariate optimization results. The table reports optimum parameters (Rmax and x) for each landscape variable, identified based on univariate optimization. The optimum spatial resolution for LULC and ecological neighborhood (scale) for density of linear features and human population density is also reported.

| Species | Variable | Rmax | x | AIC | Scale |
| --- | --- | --- | --- | --- | --- |
| JC | Human density | 1000 | 1 | -8168.29 | 5 km |
|  | LULC | 1000 | 10 | -8168.97 | 10 km |
|  | Roads | 1000 | 0.01 | -8168.34 |  |
|  | Linear density | 1000 | 2 | -8169 | 25 km |
| LPD | Human density | 1000 | 1 | -5595.08 | 10 km |
|  | LULC | 1000 | 5 | -5655.09 | 10 km |
|  | Roads | 1000 | 0.1 | -5586.22 |  |
|  | Linear density | 10 | 0.1 | -5578.62 | 0.5 km |
| SB | Human density | 1000 | 2 | -8791.33 | 10 km |
|  | LULC | 50 | 2 | -8794.12 | 0.25 km |
|  | Roads | 5 | 0.01 | -8789.5 |  |
|  | Linear density | 1000 | 2 | -8791.17 | 0.5 km |
| TIG | Human density | 50 | 2 | -9724.41 | 50 km |
|  | LULC | 100 | 2 | -10189.8 | 0.5 km |
|  | Roads | 1000 | 2 | -9522.27 |  |
|  | Linear density | 1000 | 2 | -9773.94 | 25 km and 5 km |

**Table 3.** Multivariate optimization. Optimum parameter values (Rmax and x) based on model selection using MPLE and AIC value are represented in the table. AIC null represents the AIC values for isolation by distance (null) model.

| Species | Variable | Rmax | X | AIC | AIC null |
| --- | --- | --- | --- | --- | --- |
| Jungle cat | Human density | 2 | 0.5 | -8169.03 | -8167.81 |
|  | LULC | 50 | 10 |  |  |
|  | Roads | 100 | 5 |  |  |
|  | Linear density | 1000 | 2 |  |  |
| Leopard | Human density | 50 | 1 | -5621.34 | -5571.55 |
|  | LULC | 500 | 5 |  |  |
|  | Roads | 1000 | 1 |  |  |
|  | Linear density | 2 | 1 |  |  |
| Sloth Bear | Human density | 0 | 0 | -8794.12 | -8789.47 |
|  | LULC | 50 | 2 |  |  |
|  | Roads | 0 | 0 |  |  |
|  | Linear density | 0 | 0 |  |  |
| Tiger | Human density | 0 | 0.01 | -10219.8 | -8616.83 |
|  | LULC | 100 | 2 |  |  |
|  | Roads | 1000 | 10 |  |  |
|  | Linear Density | 0 | 0.01 |  |  |

For leopards, when the landscape variables were tested separately, LULC was found to be the most important variable affecting connectivity, followed by human population density, roads and density of linear features (Table 2). The shape parameter (x>1) suggested that the relationship was nonlinear. Non-linear transformations (x >1) indicate that low resistance is offered by smaller and middle values of the variable and the resistance increases steeply with very high values. Univariate optimization identified 10 km as the optimum spatial resolution for the LULC layer, suggesting that forest cover is important at a broad scale. On combining the layers, the optimum parameters (Rmax and x) identified based on univariate optimization changed. The maximum resistance offered by human population density reduced from 1000 to 50 and by LULC from 1000 to 500. Although the maximum resistance offered by roads and density of linear features did not change, their shape parameter did (Table 3).

For sloth bears, LULC was found to be the most important variable, with an optimal resolution of 0.25 km. Human population density, roads and density of linear features also explained significant variation after controlling for Euclidean distance, when tested independently. However the magnitude of correlation and the support based on AIC scores was lower for these variables. On combining the layers, human population density, roads and density of linear features did not explain the variation in genetic differentiation over and above the impact of LULC.

The effect of landscape on genetic connectivity of tigers was found to be higher than other species, as suggested by the high model fit. LULC was the most important landscape variable explaining genetic differentiation based on univariate optimization. Density of linear features, human population density and traffic intensity on roads also explained significant variation after accounting for geographic distance (Table 2). LULC was important at a fine resolution (0.5 km), while human population density (50 km) and density of linear features (10 km) were important at larger scale for tigers. Density of linear features and human population density did not contribute to landscape resistance when all four variables were combined for multivariate optimization. Roads retained their high impact, with resistance increasing highly non-linearly with increasing traffic (Rmax=1000; x=10).

## Discussion

Understanding how anthropogenic features impact movement and connectivity of wild carnivores is important for their conservation and management. Conservation plans that incorporate data on multiple species are likely to be better than those based on a single umbrella species. Our objective was to understand how genetic variation is currently partitioned and how landscape features affect gene flow in jungle cat, leopard, sloth bear and tiger in the central Indian landscape. While the patterns of isolation by distance were in agreement with expectations based on dispersal ability and density, species’ distribution explained global genetic differentiation. Human footprint negatively impacted genetic connectivity in all four study species. The nature and scale of impact of individual landscape features was species specific. Our results provide insights into how species life-history and ecology interact with the landscape features to shape spatial genetic structure, and are important for conservation planning.

Genetic diversity in central India was high for the studied felid species and moderate for the sloth bear. Ours is the first study to estimate genetic diversity based on microsatellites in jungle cats. An earlier study using mitochondrial data on jungle cats in India had also found them to have high genetic diversity (Mukherjee et al., 2010). The only other study on leopards in the landscape (Dutta et al., 2013b) found the observed (H_o_) and expected (H_e_) heterozygosity to be higher (0.74 and 0.84 respectively), possibly due to higher sample size in their study.

Studies on tigers in the landscape have reported heterozygosity levels comparable to what this study finds (H_o_=0.52, H_e_=0.723- our study; H_o_=0.54, H_e_=0.81 (Joshi et al., 2013); H_o_=0.71, H_e_=0.75 (Yumnam et al., 2014); H_o_=0.65, H_e_=0.81 (Sharma et al., 2012)). The heterozygosity based on microsatellites was also found to be low (H_e_= 0.45) for spectacled bears that are endemic to the Andes mountain range in South America (Ruiz-García et al., 2005). The only other study on sloth bears in the landscape also found them to have moderate heterozygosity (H_o_=0.54, H_e_=0.66 (Dutta et al., 2015)).

### Dispersal ability and density impact IBD patterns

All four study species revealed significant isolation by distance (IBD). IBD relationship broke down (correlation between genetic and geographic distance was not significant) at distances beyond ∼15 km for Jungle cat, ∼40 km for sloth bear, ∼60 km for leopard and ∼140 km for tiger. Allometric scaling equations (Bowman et al., 2002) based on body size and trophic level predict the median dispersal distance in the study species to be in the same order- jungle cat (8 km), sloth bear (13 km), leopard (35 km) and tiger (85 km). Based on literature, the density at which the study species occur is also likely to follow the same order, with jungle cats likely to occur at high densities (0.4-1.5/ km^2^, Gray et al., 2016), followed by sloth bear (0.3-0.7/ km^2^, Garshelis & Smith, 1999)), leopard (0.05-0.3/ km^2^, Kalle et al., 2011; Athreya et al., 2013) and tiger (0.04-0.17/ km^2^, (Karanth & Nichols, 1998; Karanth et al., 2004; Kalle et al., 2011). The observed pattern of IBD breakdown is in accordance with what one would expect based on Wright’s genetic neighborhood size for the species (Wright, 1946). The neighborhood size depends on density of individuals and species dispersal ability and is non-linearly related to the spatial scale of autocorrelation. Although expected, such trend in breakdown of IBD has been rarely demonstrated with empirical data. It is interesting that we observe this trend despite the differential impact of landscape features on species’ geneflow. Spatial autocorrelation of genetic variation accrues over time and can remain stable over long periods (Epperson, 2005). The relatively recent landscape change may have altered the pattern of isolation by distance expected based on a more or less continuous historical distribution for all the study species. However, as observed in a recent analysis by Tucker et al (2018) for dispersal distances in terrestrial mammals, despite high impact of human footprint, body mass and dietary guild may also explain significant variation in spatial autocorrelation of genetic variation.

### Distribution shapes spatial genetic structure

The partitioning of genetic variation, as measured using F_ST_, G_ST_ and Mantel’s r, is highest for the tiger, followed by leopard and then sloth bear and jungle cat. The observed spatial genetic structure can be expected based on the distribution of the species in the central Indian landscape. Sloth bears are the most widely distributed large carnivores in central India, followed by leopard and then tiger (Jhala et al., 2015). Jungle cats are also widely distributed in central India. Their habitat association suggests a more or less continuous distribution in the landscape, although no research has been carried out. Patchily distributed populations of species are expected to have low gene flow between patches and higher impact of drift (compared to semi-continuously distributed species) that could result in high genetic differentiation (Allendorf & Luikart, 2007). This has been demonstrated in a few recent studies on plants and birds. Two comparative studies on plants find that semi-continuously or continuously distributed populations have lower genetic differentiation than patchily distributed populations (Levy et al., 2016; Llorens et al., 2017). Robin et al. (2015) studied an entire community of montane birds of the Western Ghats in India and found, with a few exceptions, highest genetic differentiation in montane-restricted species while the widespread species showed no genetic differentiation. Although comparative studies in mammals are rare, continuously distributed populations have been shown to have low genetic differentiation (Cowled et al., 2009; Côté et al., 2012; Brashear et al., 2015), unless the currently continuous population has a history of range shift or decline followed by spatial expansion (Haanes et al., 2011). Ours is the first comparative study in mammals that demonstrates a relationship between distribution and spatial genetic structure.

### Human footprint impacts all four species

Landscape genetic analysis revealed that human footprint has an impact on connectivity of the study species in the landscape. LULC was an important variable that affected genetic connectivity in all four study species. While built up areas had high impact on all the species, agriculture had a variable impact. Density of linear features had a strong negative impact on the jungle cat. Density of linear features (highway, rivers and canals) has been shown to influence genetic connectivity of roe deer (Capreolus capreolus, Coulon et al., 2006) and has also been shown to affect abundance (Beazley et al., 2004; Boulanger & Stenhouse, 2014) and occurrence (Wasserman et al., 2012) of mammals, which in-turn can impact genetic differentiation in a landscape (Weckworth et al., 2013; Llorens et al., 2017).

There has been a lot of focus on the effect of roads on wildlife populations (Van Der Ree et al., 2011; Laurance et al., 2014). Roads have been shown to reduce gene flow in bighorn sheep (Ovis Canadensis, (Epps et al., 2005)) and puma (Puma concolor, (Ernest et al., 2014)) among others. Most of these studies have been carried out on a single species (but see (Frantz et al., 2012)). Our results revealed differential impact of roads on our study species. Only roads with high intensity of traffic were found to offer high cost to tiger movement (results were consistent with individual based analysis used in this study and population based analysis carried out with a subset of tiger samples from this study in Thatte et al. (2018)). The resistance offered by roads increased gradually for leopards and jungle cats with increasing traffic on roads, with maximum resistance offered by roads being higher for leopards than jungle cats. There are several roads crisscrossing through the central Indian landscape and 70% of them have low to medium traffic intensity. These low and medium traffic roads may not impact tiger connectivity but have a high negative impact on jungle cats and leopards. Our results emphasize the need to plan and install mitigation structures on not only national and state highways with high traffic but also on smaller roads.

### Landscape variables impact species at different scales

Recent studies have suggested carrying out landscape genetic analysis at multiple spatial scales (Cushman & Landguth, 2010). Univariate optimization revealed that model fit varied for the same landscape variable (tested for the same species) at different scales (Figures S3 and S4). The optimum scale of the same landscape variable (as identified by univariate optimization) was different for the study species. For example, optimum scale of the human population density layer was 50km for tigers, while it was 10 km for leopard and sloth bear and 5 km for jungle cats.

### Low genetic differentiation and landscape genetics analysis

Simulation based studies have found that landscape effects, inferred based on landscape genetic analyses, are most reliable when genetic differentiation is high (Zeller et al., 2016; Shirk et al., 2017) and/or the resistance offered by the landscape is high (Jaquiery et al., 2011). We found low genetic differentiation in jungle cats and sloth bears. Both these species are likely to have high effective population sizes, given their wide distribution in central India. When population size in habitat fragments is high, only a few successful dispersal events are required to counter genetic differentiation due to drift (Mills & Allendorf, 1996). Kekkonen (2011) investigated genetic differentiation in house sparrows in a continuously distributed population. Although dispersal ability of sparrows is low (only 10% of ringed birds were found to disperse >16km), authors found negligible genetic differentiation across Finland (covering an area of 400 x 800 km). However, genetic differentiation was high between populations in Finland and Sweden that were separated by 40 km of open sea, a biogeographic barrier. In cases where there is a lack of discrete population boundaries and the genetic differentiation is low, it is a challenge to identify effect of landscape features on gene flow. The model fit in case of sloth bears and jungle cats was also poor. Poor model fit may not imply lack of landscape effects on connectivity in these species, but the inability of landscape genetics methods to detect those. However, low model fit along with low genetic differentiation may suggest a lack of absolute barriers in the landscape for the species. Our result highlight the need to combine different data sources and develop methods complementary to landscape genetics to understand the impact of landscape features on species with low genetic differentiation.

### Interactions between species life-history, ecology and landscape structure

We see interesting genetic patterns in our data that are influenced by both species morphology and ecology. We found dispersal ability impacted spatial autocorrelation between genetic and geographic distance in the study species, while distribution seems to explain the genetic differentiation pattern. Often considered in isolation in comparative studies, species life-history and ecological characteristics are likely to interact with each other and with landscape characteristics to shape spatial genetic structure. All three variables-habitat association, dispersal ability and distribution predicted that jungle cats would show the lowest genetic differentiation among the study species and our results corroborated these expectations. Both tigers and sloth bears were expected to show high genetic differentiation based on two of the three variables. While tiger had the highest genetic structure among the study species and a strong effect of landscape features, sloth bears had low genetic differentiation and poor model fit in landscape genetic analysis. Sloth bears are likely to have a high effective population size owing to their high densities (Garshelis & Smith, 1999; Ratnayeke et al., 2007), ability to survive in small habitat patches (Bargali et al., 2012) and wide distribution in central India (Jhala et al., 2015). Hence, despite low to moderate dispersal ability, they are likely to experience low genetic drift resulting in low genetic differentiation. Additionally, sloth bear demographic decline has likely been gradual over the last few decades (Dharaiya et al., 2016). On the other hand, models of tiger demographic history suggest they went through a population bottleneck ∼200 years ago and lost 90% of the population (Mondol et al., 2013). The high genetic structure observed in tigers, despite high dispersal ability, could also be due to a combination of demographic history of population decline and their current restricted distribution.

The observed patterns of connectivity in landscape genetic studies are likely to be a result of multiple interacting processes that influence gene flow and genetic drift. There is a dearth of conceptual and theoretical research investigating the interactions between the different variables including dispersal ability, habitat specialization, distribution and demographic history along with landscape characteristics (area and configuration) and their effect on gene flow and genetic drift (Balkenhol et al., 2016).

### Conservation implications

Conservation tends to focus on large mammals in protected areas. While this is important, it may not be an effective conservation strategy for widely distributed and genetically well connected populations of species like the jungle cat and sloth bear. For such species, conservation value of non-protected areas and small habitat fragments might be high despite being embedded in a human dominated landscape. Along with delineating corridors between protected areas that are critical for persistence of species like the tiger (Qureshi et al., 2014; Thatte et al., 2018a), landscape scale conservation strategies also need to factor in the requirements of species that are abundant outside protected areas and currently well connected. Considering the rapid rate of deforestation and anthropogenic development in forest and non-forest areas, it is important to realize the conservation value of modified habitats (Pardini et al., 2009) and non-protected areas for conserving multi-species connectivity. Mitigation measures on linear infrastructure also need to consider the needs of multiple species. While installing mitigation measures on wide roads with high traffic might be sufficient to maintain tiger connectivity, it may not be enough for leopards. For jungle cats, point corridors across roads may not be enough as it is the density of linear features that has a strong negative impact on its connectivity. Given the differential impacts, we acknowledge that conservation planning to maintain multi-species connectivity can be a challenge. Any planning approach that aims to conserve multiple species will be a compromise between what is best for each species individually and what is optimum and feasible when all species are considered together (Early & Thomas, 2007). Indeed, very few studies have tried to incorporate data on multiple species that differ in their connectivity pattern and dispersal ability into conservation planning exercises (but see Early & Thomas 2007; Magris et al. 2016; Dudaniec et al. 2016).

### Conclusion

Landscapes of high conservation value, especially in the tropics, have been changing rapidly due to fragmentation and degradation of habitat (Crooks et al., 2011; Elmhagen et al., 2015; Newbold et al., 2015). Understanding the impact of fragmentation on multiple species and identifying strategies to maintain connectivity are critical to counter the negative impacts of fragmentation. By investigating how landscape features impact connectivity, studies like ours can help towards a proactive approach to conservation that can anticipate and prevent future species declines. Species persistence being one of the key goals of conservation planning (Pereira et al., 2010), a better conceptual understanding of how species interact with their landscape is imperative. Multi-species studies are also useful to gain insights into understanding how species traits interact with landscape characteristics to shape spatial genetic structure, linking ecological and evolutionary processes. Using a combination of empirical data on multiple species and spatially explicit simulations, future studies can work towards a conceptual and theoretical understanding of the interacting processes shaping spatial distribution of genetic variation.

## Data accessibility statement

Genotypes of final set of individuals used for analysis will be uploaded on Dryad repository.

