## Supplementary material for "Human footprint differentially impacts genetic connectivity of four wide-ranging mammals in a fragmented landscape"

Figure S1. Landscape variables used for resistance optimization

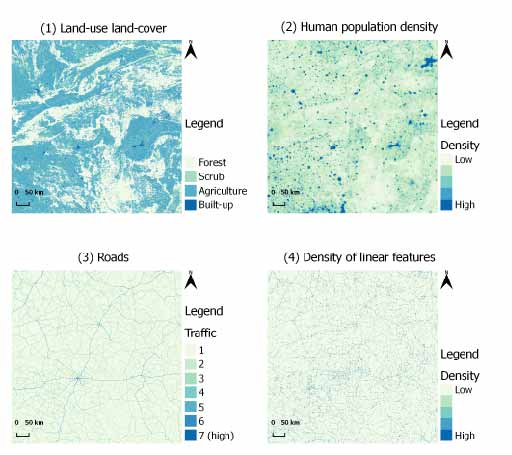

Figure S2. Six curves used to transform landscape variable values into resistance values. The curves are based on the function R=1+Rmax((V-1)/(Vmax-1))^x^, where R is the resistance, Rmax is the maximum resistance (varied at six levels in the paper, fixed here in the figure at 1000), V is the variable value at each pixel, Vmax is the maximum variable value and x (varied at six levels 0.001, 0.1, 1, 2, 5, 10) determines the shape of the curve.

Shape transformations

Cost/ resistance

| Density of linear features | 0.01 | 0.1 | 0.5 | 1 | 2 | 5 | 10 |
| --- | --- | --- | --- | --- | --- | --- | --- |
| 1 | 1 | 1 | 1 | 1 | 1 | 1 | 1 |
| 1.30572 | 881.386 | 280.724 | 2.71257 | 1.00293 | 1 | 1 | 1 |
| 1.58206 | 887.073 | 299.328 | 3.36302 | 1.00558 | 1 | 1 | 1 |
| 2.02256 | 892.08 | 316.621 | 4.13205 | 1.00981 | 1 | 1 | 1 |
| 2.23248 | 893.745 | 322.57 | 4.43855 | 1.01182 | 1 | 1 | 1 |
| 4.27555 | 902.514 | 355.589 | 6.60567 | 1.03142 | 1 | 1 | 1 |
| 4.48294 | 903.068 | 357.773 | 6.7804 | 1.03341 | 1 | 1 | 1 |
| 6.5381 | 907.261 | 374.709 | 8.28896 | 1.05313 | 1 | 1 | 1 |
| 7.35503 | 908.509 | 379.886 | 8.80808 | 1.06097 | 1 | 1 | 1 |
| 8.69286 | 910.244 | 387.194 | 9.59071 | 1.0738 | 1.00001 | 1 | 1 |
| 8.79263 | 910.361 | 387.692 | 9.64624 | 1.07476 | 1.00001 | 1 | 1 |
| 11.329 | 912.927 | 398.743 | 10.9544 | 1.09909 | 1.00001 | 1 | 1 |
| 12.6978 | 914.063 | 403.724 | 11.5935 | 1.11222 | 1.00001 | 1 | 1 |
| 13.2294 | 914.469 | 405.518 | 11.8315 | 1.11732 | 1.00001 | 1 | 1 |
| 13.8058 | 914.89 | 407.385 | 12.0838 | 1.12285 | 1.00002 | 1 | 1 |
| 14.5275 | 915.391 | 409.619 | 12.3918 | 1.12977 | 1.00002 | 1 | 1 |
| 16.074 | 916.381 | 414.066 | 13.0254 | 1.14461 | 1.00002 | 1 | 1 |
| 20.9759 | 918.962 | 425.862 | 14.8433 | 1.19164 | 1.00004 | 1 | 1 |
| 21.4883 | 919.195 | 426.939 | 15.0197 | 1.19655 | 1.00004 | 1 | 1 |
| 22.9929 | 919.846 | 429.968 | 15.5254 | 1.21099 | 1.00004 | 1 | 1 |
| 23.165 | 919.917 | 430.303 | 15.5821 | 1.21264 | 1.00005 | 1 | 1 |
| 23.4275 | 920.025 | 430.809 | 15.6682 | 1.21515 | 1.00005 | 1 | 1 |
| 24.1746 | 920.327 | 432.219 | 15.9105 | 1.22232 | 1.00005 | 1 | 1 |
| 25.5443 | 920.855 | 434.703 | 16.3448 | 1.23546 | 1.00006 | 1 | 1 |
| 27.0603 | 921.406 | 437.31 | 16.8116 | 1.25001 | 1.00006 | 1 | 1 |
| 27.101 | 921.42 | 437.378 | 16.8239 | 1.2504 | 1.00006 | 1 | 1 |
| 28.2339 | 921.812 | 439.236 | 17.1637 | 1.26126 | 1.00007 | 1 | 1 |
| 28.6754 | 921.96 | 439.941 | 17.2942 | 1.2655 | 1.00007 | 1 | 1 |
| 29.587 | 922.258 | 441.366 | 17.5603 | 1.27424 | 1.00008 | 1 | 1 |
| 31.5697 | 922.876 | 444.329 | 18.125 | 1.29327 | 1.00009 | 1 | 1 |
| 32.3215 | 923.1 | 445.407 | 18.3343 | 1.30048 | 1.00009 | 1 | 1 |
| 32.7013 | 923.211 | 445.943 | 18.4391 | 1.30412 | 1.00009 | 1 | 1 |
| 32.7148 | 923.215 | 445.962 | 18.4428 | 1.30425 | 1.00009 | 1 | 1 |
| 33.377 | 923.406 | 446.883 | 18.6239 | 1.3106 | 1.0001 | 1 | 1 |
| 33.8826 | 923.549 | 447.574 | 18.761 | 1.31545 | 1.0001 | 1 | 1 |
| 35.4859 | 923.988 | 449.705 | 19.1889 | 1.33084 | 1.00011 | 1 | 1 |
| 38.6663 | 924.803 | 453.681 | 20.0091 | 1.36135 | 1.00013 | 1 | 1 |
| 40.4445 | 925.229 | 455.774 | 20.4526 | 1.37841 | 1.00014 | 1 | 1 |
| 41.4675 | 925.466 | 456.94 | 20.7033 | 1.38822 | 1.00015 | 1 | 1 |
| 41.8488 | 925.552 | 457.368 | 20.7959 | 1.39188 | 1.00015 | 1 | 1 |
| 42.1694 | 925.625 | 457.725 | 20.8734 | 1.39495 | 1.00016 | 1 | 1 |
| 42.6466 | 925.731 | 458.251 | 20.9883 | 1.39953 | 1.00016 | 1 | 1 |
| 42.8299 | 925.772 | 458.452 | 21.0322 | 1.40129 | 1.00016 | 1 | 1 |
| 43.1762 | 925.848 | 458.829 | 21.1149 | 1.40461 | 1.00016 | 1 | 1 |
| 44.8934 | 926.217 | 460.66 | 21.5203 | 1.42108 | 1.00018 | 1 | 1 |
| 45.1775 | 926.277 | 460.957 | 21.5867 | 1.42381 | 1.00018 | 1 | 1 |
| 46.1402 | 926.476 | 461.95 | 21.8098 | 1.43305 | 1.00019 | 1 | 1 |
| 47.598 | 926.771 | 463.417 | 22.1431 | 1.44703 | 1.0002 | 1 | 1 |
| 48.1468 | 926.879 | 463.959 | 22.2673 | 1.4523 | 1.0002 | 1 | 1 |
| 49.1361 | 927.071 | 464.921 | 22.4892 | 1.46179 | 1.00021 | 1 | 1 |
| 49.3516 | 927.113 | 465.128 | 22.5373 | 1.46385 | 1.00022 | 1 | 1 |
| 49.4108 | 927.124 | 465.185 | 22.5505 | 1.46442 | 1.00022 | 1 | 1 |
| 49.6314 | 927.166 | 465.396 | 22.5995 | 1.46654 | 1.00022 | 1 | 1 |
| 49.9691 | 927.23 | 465.718 | 22.6744 | 1.46978 | 1.00022 | 1 | 1 |
| 51.648 | 927.542 | 467.287 | 23.0428 | 1.48588 | 1.00024 | 1 | 1 |
| 53.4997 | 927.875 | 468.964 | 23.4421 | 1.50365 | 1.00025 | 1 | 1 |
| 53.7895 | 927.926 | 469.222 | 23.504 | 1.50643 | 1.00026 | 1 | 1 |
| 54.481 | 928.047 | 469.832 | 23.6509 | 1.51306 | 1.00026 | 1 | 1 |
| 55.1553 | 928.163 | 470.419 | 23.7932 | 1.51953 | 1.00027 | 1 | 1 |
| 55.3174 | 928.191 | 470.56 | 23.8273 | 1.52109 | 1.00027 | 1 | 1 |
| 57.4173 | 928.543 | 472.344 | 24.2644 | 1.54123 | 1.00029 | 1 | 1 |
| 58.5941 | 928.734 | 473.318 | 24.5058 | 1.55252 | 1.00031 | 1 | 1 |
| 59.1411 | 928.822 | 473.765 | 24.6171 | 1.55777 | 1.00031 | 1 | 1 |
| 61.4625 | 929.185 | 475.62 | 25.084 | 1.58004 | 1.00034 | 1 | 1 |
| 62.3041 | 929.313 | 476.276 | 25.251 | 1.58811 | 1.00035 | 1 | 1 |
| 63.6668 | 929.517 | 477.322 | 25.5191 | 1.60118 | 1.00036 | 1 | 1 |
| 63.8351 | 929.542 | 477.45 | 25.552 | 1.6028 | 1.00036 | 1 | 1 |
| 64.6602 | 929.664 | 478.072 | 25.7126 | 1.61071 | 1.00037 | 1 | 1 |
| 67.1994 | 930.027 | 479.941 | 26.2007 | 1.63507 | 1.0004 | 1 | 1 |
| 67.5344 | 930.074 | 480.183 | 26.2644 | 1.63829 | 1.00041 | 1 | 1 |
| 68.1314 | 930.157 | 480.612 | 26.3775 | 1.64402 | 1.00041 | 1 | 1 |
| 70.1597 | 930.433 | 482.041 | 26.758 | 1.66347 | 1.00044 | 1 | 1 |
| 71.2321 | 930.576 | 482.782 | 26.9569 | 1.67376 | 1.00045 | 1 | 1 |
| 71.2435 | 930.578 | 482.79 | 26.959 | 1.67387 | 1.00045 | 1 | 1 |
| 71.8978 | 930.664 | 483.237 | 27.0796 | 1.68015 | 1.00046 | 1 | 1 |
| 72.4554 | 930.737 | 483.615 | 27.182 | 1.6855 | 1.00047 | 1 | 1 |
| 73.8182 | 930.913 | 484.527 | 27.4305 | 1.69857 | 1.00049 | 1 | 1 |
| 74.0888 | 930.947 | 484.707 | 27.4796 | 1.70117 | 1.00049 | 1 | 1 |
| 75.0457 | 931.068 | 485.336 | 27.6523 | 1.71035 | 1.0005 | 1 | 1 |
| 78.4897 | 931.491 | 487.543 | 28.2651 | 1.74339 | 1.00055 | 1 | 1 |
| 79.0006 | 931.552 | 487.863 | 28.3548 | 1.74829 | 1.00056 | 1 | 1 |
| 80.1166 | 931.684 | 488.555 | 28.5498 | 1.75899 | 1.00058 | 1 | 1 |
| 80.4119 | 931.719 | 488.737 | 28.6012 | 1.76183 | 1.00058 | 1 | 1 |
| 80.8038 | 931.765 | 488.977 | 28.6692 | 1.76559 | 1.00059 | 1 | 1 |
| 81.3209 | 931.825 | 489.292 | 28.7587 | 1.77055 | 1.00059 | 1 | 1 |
| 81.7202 | 931.871 | 489.535 | 28.8276 | 1.77438 | 1.0006 | 1 | 1 |
| 84.2965 | 932.164 | 491.072 | 29.2682 | 1.79909 | 1.00064 | 1 | 1 |
| 84.9507 | 932.236 | 491.455 | 29.379 | 1.80537 | 1.00065 | 1 | 1 |
| 85.3898 | 932.285 | 491.711 | 29.4531 | 1.80958 | 1.00066 | 1 | 1 |
| 86.222 | 932.376 | 492.193 | 29.5931 | 1.81756 | 1.00067 | 1 | 1 |
| 87.6354 | 932.53 | 493.002 | 29.8292 | 1.83112 | 1.00069 | 1 | 1 |
| 88.3608 | 932.607 | 493.412 | 29.9497 | 1.83808 | 1.0007 | 1 | 1 |
| 89.0564 | 932.681 | 493.803 | 30.0647 | 1.84476 | 1.00071 | 1 | 1 |
| 89.1105 | 932.687 | 493.833 | 30.0736 | 1.84527 | 1.00071 | 1 | 1 |
| 89.8153 | 932.761 | 494.226 | 30.1897 | 1.85204 | 1.00073 | 1 | 1 |
| 90.4574 | 932.828 | 494.581 | 30.295 | 1.8582 | 1.00074 | 1 | 1 |
| 92.6607 | 933.055 | 495.784 | 30.6536 | 1.87933 | 1.00077 | 1 | 1 |
| 92.9479 | 933.084 | 495.939 | 30.7 | 1.88209 | 1.00078 | 1 | 1 |
| 93.8493 | 933.175 | 496.422 | 30.8452 | 1.89074 | 1.00079 | 1 | 1 |
| 96.2505 | 933.413 | 497.688 | 31.2287 | 1.91377 | 1.00083 | 1 | 1 |
| 96.479 | 933.435 | 497.807 | 31.2649 | 1.91596 | 1.00084 | 1 | 1 |
| 96.5928 | 933.447 | 497.866 | 31.2829 | 1.91706 | 1.00084 | 1 | 1 |
| 98.0157 | 933.584 | 498.601 | 31.5075 | 1.93071 | 1.00087 | 1 | 1 |
| 98.4676 | 933.628 | 498.832 | 31.5784 | 1.93504 | 1.00087 | 1 | 1 |
| 98.6147 | 933.642 | 498.907 | 31.6015 | 1.93645 | 1.00088 | 1 | 1 |
| 98.6249 | 933.643 | 498.913 | 31.6031 | 1.93655 | 1.00088 | 1 | 1 |
| 101.663 | 933.929 | 500.441 | 32.0757 | 1.9657 | 1.00093 | 1 | 1 |
| 102.556 | 934.011 | 500.882 | 32.2132 | 1.97427 | 1.00095 | 1 | 1 |
| 103.665 | 934.112 | 501.425 | 32.3831 | 1.9849 | 1.00097 | 1 | 1 |
| 106.365 | 934.355 | 502.726 | 32.7932 | 2.01081 | 1.00102 | 1 | 1 |
| 111.287 | 934.781 | 505.022 | 33.5273 | 2.05802 | 1.00112 | 1 | 1 |
| 113.938 | 935.003 | 506.221 | 33.9159 | 2.08345 | 1.00117 | 1 | 1 |
| 115.232 | 935.109 | 506.796 | 34.1038 | 2.09586 | 1.0012 | 1 | 1 |
| 115.462 | 935.128 | 506.898 | 34.1371 | 2.09807 | 1.00121 | 1 | 1 |
| 120.276 | 935.513 | 508.987 | 34.8269 | 2.14426 | 1.00131 | 1 | 1 |
| 122.449 | 935.681 | 509.905 | 35.1336 | 2.1651 | 1.00136 | 1 | 1 |
| 122.582 | 935.692 | 509.96 | 35.1523 | 2.16638 | 1.00136 | 1 | 1 |
| 123.723 | 935.779 | 510.436 | 35.3121 | 2.17732 | 1.00139 | 1 | 1 |
| 127.259 | 936.045 | 511.885 | 35.8029 | 2.21124 | 1.00147 | 1 | 1 |
| 129.109 | 936.181 | 512.629 | 36.057 | 2.229 | 1.00151 | 1 | 1 |
| 130.071 | 936.251 | 513.012 | 36.1885 | 2.23823 | 1.00153 | 1 | 1 |
| 130.122 | 936.254 | 513.032 | 36.1953 | 2.23871 | 1.00153 | 1 | 1 |
| 130.46 | 936.279 | 513.166 | 36.2415 | 2.24196 | 1.00154 | 1 | 1 |
| 131.298 | 936.339 | 513.496 | 36.3552 | 2.24999 | 1.00156 | 1 | 1 |
| 134.774 | 936.585 | 514.848 | 36.8238 | 2.28335 | 1.00165 | 1 | 1 |
| 135.502 | 936.636 | 515.127 | 36.9212 | 2.29033 | 1.00166 | 1 | 1 |
| 135.773 | 936.655 | 515.23 | 36.9573 | 2.29293 | 1.00167 | 1 | 1 |
| 137.274 | 936.759 | 515.8 | 37.157 | 2.30733 | 1.00171 | 1 | 1 |
| 139.015 | 936.877 | 516.454 | 37.3872 | 2.32403 | 1.00175 | 1 | 1 |
| 139.367 | 936.901 | 516.585 | 37.4335 | 2.3274 | 1.00176 | 1 | 1 |
| 141.673 | 937.056 | 517.438 | 37.7359 | 2.34953 | 1.00182 | 1 | 1 |
| 143.084 | 937.149 | 517.954 | 37.9196 | 2.36306 | 1.00186 | 1 | 1 |
| 144.121 | 937.217 | 518.33 | 38.0541 | 2.37301 | 1.00189 | 1 | 1 |
| 147.161 | 937.414 | 519.418 | 38.4456 | 2.40217 | 1.00197 | 1 | 1 |
| 148.228 | 937.482 | 519.795 | 38.582 | 2.41241 | 1.00199 | 1 | 1 |
| 148.514 | 937.501 | 519.896 | 38.6186 | 2.41516 | 1.002 | 1 | 1 |
| 148.598 | 937.506 | 519.926 | 38.6292 | 2.41596 | 1.002 | 1 | 1 |
| 148.661 | 937.51 | 519.948 | 38.6373 | 2.41657 | 1.00201 | 1 | 1 |
| 149.384 | 937.556 | 520.201 | 38.7294 | 2.4235 | 1.00203 | 1 | 1 |
| 149.455 | 937.56 | 520.226 | 38.7384 | 2.42419 | 1.00203 | 1 | 1 |
| 150.596 | 937.632 | 520.624 | 38.8831 | 2.43513 | 1.00206 | 1 | 1 |
| 150.927 | 937.653 | 520.739 | 38.925 | 2.4383 | 1.00207 | 1 | 1 |
| 152.898 | 937.775 | 521.418 | 39.1734 | 2.45721 | 1.00212 | 1 | 1 |
| 154.191 | 937.854 | 521.859 | 39.3355 | 2.46961 | 1.00216 | 1 | 1 |
| 155.327 | 937.923 | 522.244 | 39.4774 | 2.48051 | 1.00219 | 1 | 1 |
| 159.69 | 938.185 | 523.699 | 40.0176 | 2.52237 | 1.00232 | 1 | 1 |
| 162.281 | 938.336 | 524.546 | 40.3347 | 2.54722 | 1.00239 | 1 | 1 |
| 165.122 | 938.5 | 525.462 | 40.6797 | 2.57448 | 1.00248 | 1 | 1 |
| 166.138 | 938.558 | 525.785 | 40.8023 | 2.58422 | 1.00251 | 1 | 1 |
| 167.305 | 938.624 | 526.155 | 40.9428 | 2.59542 | 1.00255 | 1 | 1 |
| 167.821 | 938.653 | 526.318 | 41.0047 | 2.60037 | 1.00256 | 1 | 1 |
| 168.43 | 938.687 | 526.509 | 41.0777 | 2.60622 | 1.00258 | 1 | 1 |
| 168.535 | 938.693 | 526.542 | 41.0901 | 2.60722 | 1.00258 | 1 | 1 |
| 170.086 | 938.78 | 527.027 | 41.2754 | 2.62211 | 1.00263 | 1 | 1 |
| 170.403 | 938.797 | 527.125 | 41.313 | 2.62514 | 1.00264 | 1 | 1 |
| 171.846 | 938.877 | 527.572 | 41.4844 | 2.63899 | 1.00269 | 1 | 1 |
| 172.596 | 938.918 | 527.802 | 41.5731 | 2.64618 | 1.00271 | 1 | 1 |
| 173.712 | 938.979 | 528.144 | 41.7049 | 2.65689 | 1.00275 | 1 | 1 |
| 173.976 | 938.993 | 528.224 | 41.7359 | 2.65942 | 1.00275 | 1 | 1 |
| 177.596 | 939.187 | 529.318 | 42.16 | 2.69415 | 1.00287 | 1 | 1 |
| 177.783 | 939.197 | 529.374 | 42.1818 | 2.69594 | 1.00288 | 1 | 1 |
| 178.658 | 939.243 | 529.635 | 42.2836 | 2.70434 | 1.0029 | 1 | 1 |
| 178.978 | 939.26 | 529.73 | 42.3207 | 2.7074 | 1.00292 | 1 | 1 |
| 179.272 | 939.276 | 529.817 | 42.3548 | 2.71022 | 1.00292 | 1 | 1 |
| 179.537 | 939.29 | 529.895 | 42.3856 | 2.71277 | 1.00293 | 1 | 1 |
| 184.501 | 939.547 | 531.348 | 42.957 | 2.76039 | 1.0031 | 1 | 1 |
| 186.389 | 939.643 | 531.891 | 43.1723 | 2.7785 | 1.00316 | 1 | 1 |
| 187.887 | 939.719 | 532.319 | 43.3424 | 2.79288 | 1.00321 | 1 | 1 |
| 188.102 | 939.73 | 532.38 | 43.3667 | 2.79494 | 1.00322 | 1 | 1 |
| 189.407 | 939.795 | 532.749 | 43.5142 | 2.80746 | 1.00327 | 1 | 1 |
| 189.485 | 939.799 | 532.771 | 43.523 | 2.80821 | 1.00327 | 1 | 1 |
| 191.336 | 939.89 | 533.291 | 43.7313 | 2.82596 | 1.00333 | 1 | 1 |
| 191.402 | 939.894 | 533.31 | 43.7387 | 2.8266 | 1.00334 | 1 | 1 |
| 191.609 | 939.904 | 533.367 | 43.7619 | 2.82858 | 1.00334 | 1 | 1 |
| 195.622 | 940.1 | 534.478 | 44.2097 | 2.86708 | 1.00349 | 1 | 1 |
| 195.907 | 940.113 | 534.556 | 44.2413 | 2.86981 | 1.0035 | 1 | 1 |
| 197.415 | 940.186 | 534.967 | 44.4083 | 2.88428 | 1.00355 | 1 | 1 |
| 197.747 | 940.202 | 535.057 | 44.4449 | 2.88746 | 1.00356 | 1 | 1 |
| 198.059 | 940.216 | 535.142 | 44.4794 | 2.89045 | 1.00357 | 1 | 1 |
| 198.107 | 940.219 | 535.155 | 44.4847 | 2.89092 | 1.00358 | 1 | 1 |
| 198.109 | 940.219 | 535.156 | 44.4849 | 2.89094 | 1.00358 | 1 | 1 |
| 198.335 | 940.23 | 535.217 | 44.5099 | 2.89311 | 1.00358 | 1 | 1 |
| 199.254 | 940.273 | 535.465 | 44.611 | 2.90192 | 1.00362 | 1 | 1 |
| 200.484 | 940.331 | 535.796 | 44.7461 | 2.91372 | 1.00366 | 1 | 1 |
| 202.395 | 940.421 | 536.306 | 44.9551 | 2.93205 | 1.00373 | 1 | 1 |
| 203.295 | 940.463 | 536.544 | 45.0532 | 2.94068 | 1.00377 | 1 | 1 |
| 204.347 | 940.511 | 536.822 | 45.1676 | 2.95078 | 1.00381 | 1 | 1 |
| 205.892 | 940.583 | 537.228 | 45.3351 | 2.9656 | 1.00386 | 1 | 1 |
| 206.65 | 940.617 | 537.426 | 45.4171 | 2.97288 | 1.00389 | 1 | 1 |
| 209.692 | 940.755 | 538.214 | 45.7443 | 3.00206 | 1.00401 | 1 | 1 |
| 210.214 | 940.779 | 538.349 | 45.8003 | 3.00707 | 1.00403 | 1 | 1 |
| 212.174 | 940.866 | 538.85 | 46.0096 | 3.02587 | 1.0041 | 1 | 1 |
| 215.342 | 941.006 | 539.651 | 46.346 | 3.05626 | 1.00423 | 1 | 1 |
| 215.452 | 941.011 | 539.679 | 46.3576 | 3.05731 | 1.00423 | 1 | 1 |
| 215.733 | 941.023 | 539.75 | 46.3874 | 3.06001 | 1.00424 | 1 | 1 |
| 218.208 | 941.131 | 540.367 | 46.6482 | 3.08376 | 1.00434 | 1 | 1 |
| 218.361 | 941.138 | 540.405 | 46.6642 | 3.08522 | 1.00435 | 1 | 1 |
| 218.753 | 941.155 | 540.503 | 46.7054 | 3.08898 | 1.00436 | 1 | 1 |
| 218.857 | 941.159 | 540.528 | 46.7163 | 3.08998 | 1.00437 | 1 | 1 |
| 218.945 | 941.163 | 540.55 | 46.7255 | 3.09082 | 1.00437 | 1 | 1 |

Figure S3. Relationship between spatial resolution and model fit. Data in different colours represents three different landscape variables. Lowest AIC and Highest partial Mantel’s r were the best models chosen based on the respective model selection methods.

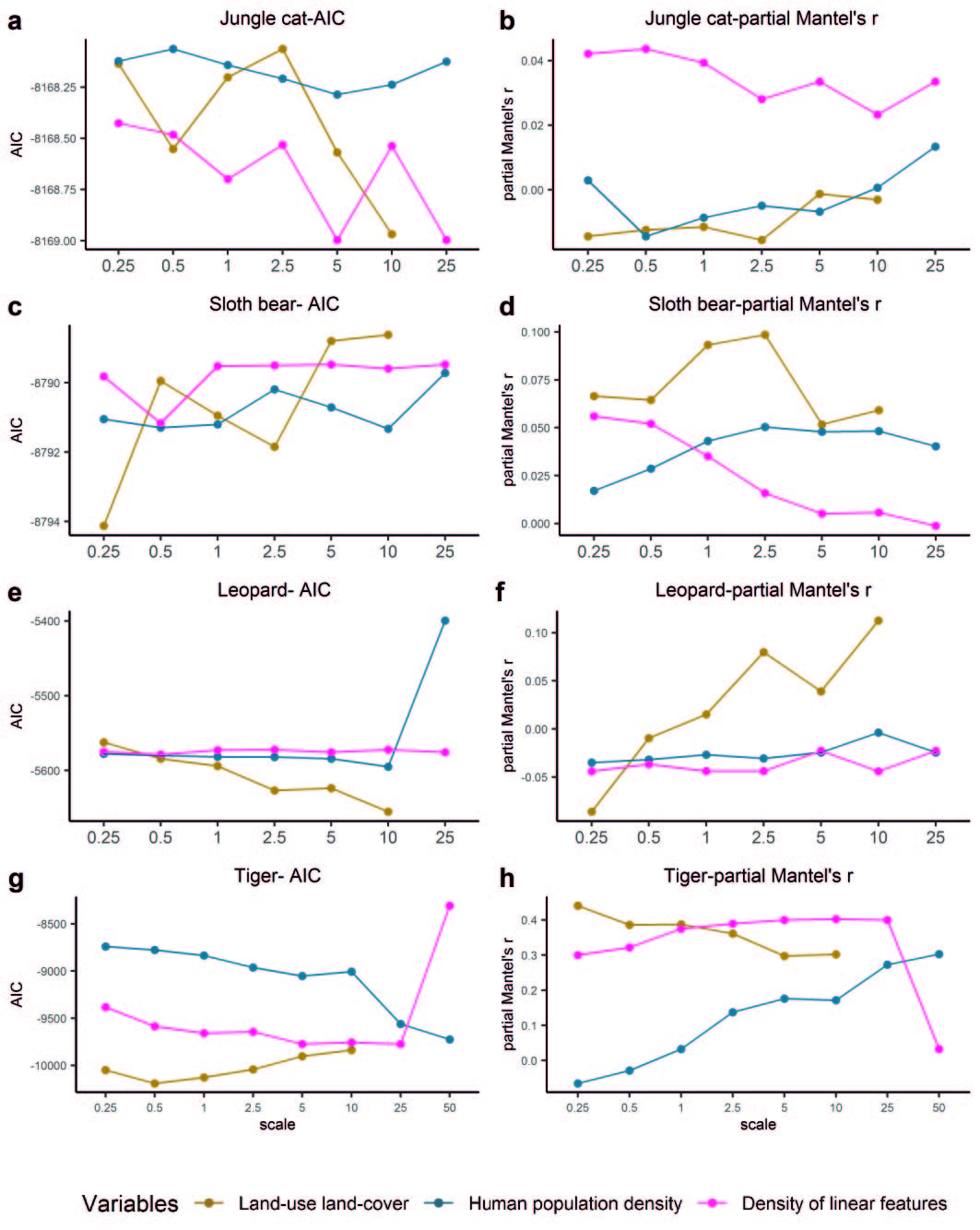

Figure S4. Performance of each variable based on model fit across spatial resolutions. For each variable on the x axis, different colours represent different spatial resolutions. Lowest AIC and Highest partial Mantel’s r were the best models chosen based on the respective model selection methods.

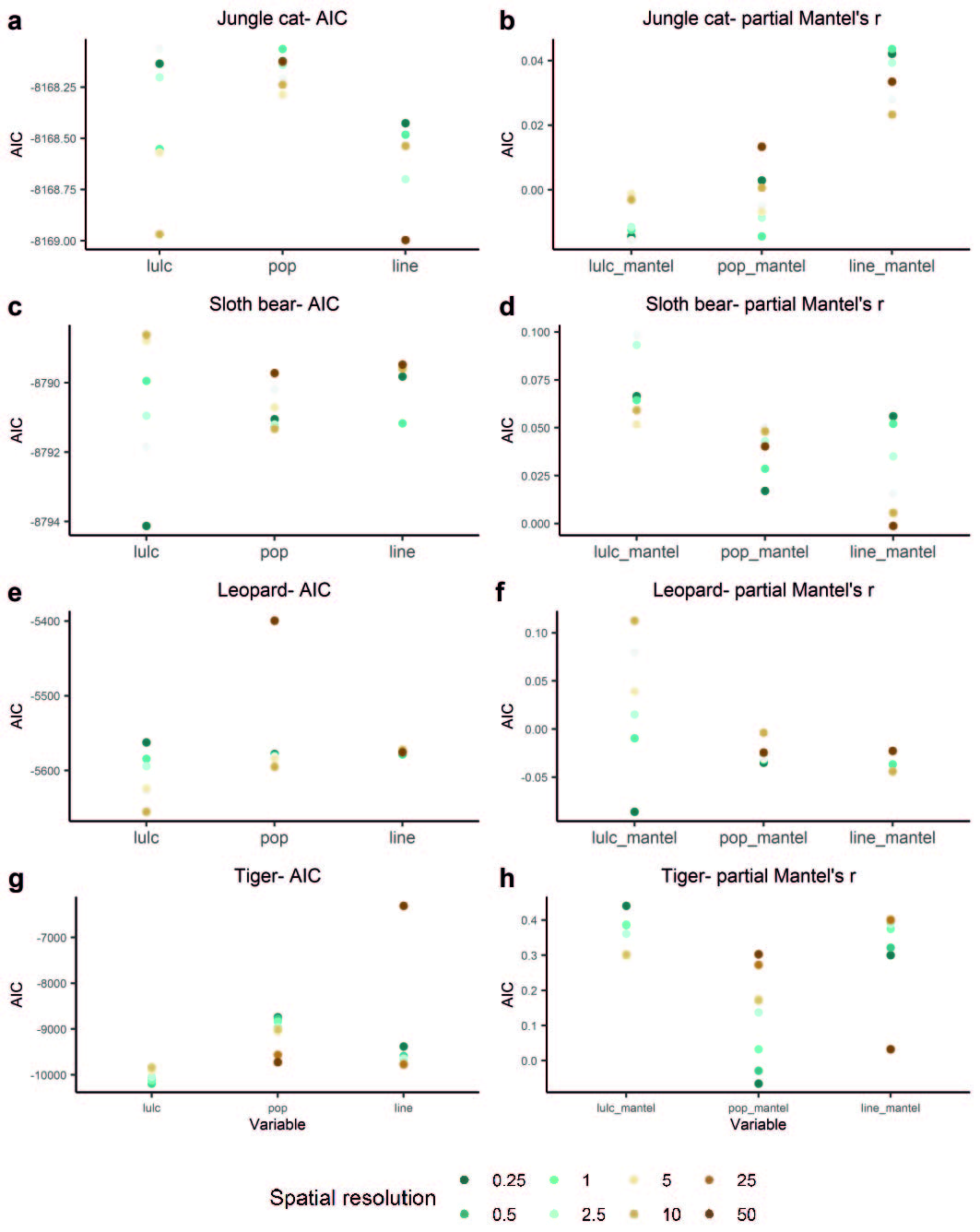

Table S1. Correlation between landscape variables

|  | **Land-use land-cover** | **Human population density** | **Roads** | **Density of linear features** |
| --- | --- | --- | --- | --- |
| **Land-use land-cover** | 1 |  |  |  |
| **Human population density** | 0.13447736 | 1 |  |  |
| **Roads** | 0.49993435 | 0.03232793 | 1 |  |
| **Density of linear features** | 0.03870765 | 0.05604515 | 0.03142308 | 1 |
